# Distinct CED-10/Rac1 Domains Confer Context-Specific Functions in Neuronal Development

**DOI:** 10.1101/341412

**Authors:** Steffen Nørgaard, Shuer Deng, Wei Cao, Roger Pocock

**Affiliations:** Development and Stem Cells Program, Monash Biomedicine Discovery Institute and Department of Anatomy and Developmental Biology, Monash University, Melbourne, Victoria 3800, Australia.; Biotech Research and Innovation Centre, University of Copenhagen, Ole Maaløes Vej 5, Copenhagen, Denmark.

## Abstract

Rac GTPases act as master switches to coordinate multiple interweaved signaling pathways. A major function for Rac GTPases is to control neurite development by influencing downstream effector molecules and pathways. In *Caenorhabditis elegans*, the Rac proteins CED-10, RAC-2 and MIG-2 act in parallel to control axon outgrowth and guidance. Here, we have identified a single glycine residue in the CED-10/Rac1 Switch 1 region that confers a non-redundant function in axon outgrowth but not guidance. Mutation of this glycine to glutamic acid (G30E) reduces GTP binding and inhibits axon outgrowth but does not affect other canonical CED-10 functions. This demonstrates previously unappreciated domain-specific functions within the CED-10 protein. Further, we reveal that when CED-10 function is diminished, the adaptor protein NAB-1 (Neurabin) and its interacting partner SYD-1 (Rho-GAP-like protein) can act as inhibitors of axon outgrowth. Together, we reveal that specific domains and residues within Rac GTPases can confer context-dependent functions during animal development.

---

Correct axonal projection is essential for developing a functional nervous system. The *Caenorhabditis elegans* ventral nerve cord (VNC) contains two fascicles housing multiple neurons that project axons along the length of the animal^1^. During development, the extending tips of VNC axons (growth cones) facilitate faithful navigation of their environment by interpreting outgrowth and guidance signals^2^^-^^10^. Growth cones assimilate molecular information from neighbouring cells and the extracellular environment through ligand-receptor interactions. Receptors then regulate complex and interconnected intracellular signaling cascades that ultimately control cytoskeletal dynamics^11, 12^. These signalling mechanisms are tightly orchestrated, both temporally and spatially, to ensure precise nervous system development.

Rho GTPase family members (Rho, Rac and Cdc42) are key regulators of actin cytoskeletal dynamics^13^, and are especially recognized for their role in regulating axon outgrowth and guidance^14, 15^. The three *C. elegans* Rac GTPases, CED-10, MIG-2 and RAC-2 have overlapping functions in the context of axon outgrowth and guidance^16^. As such, classical single mutants in any of the *C. elegans* Rac GTPases result in minor axonal defects with compound mutations having synergistic effects on outgrowth and guidance^16^. A major function for Rac GTPases is to coordinate actin filament networks at the growth cone of an extending axon^17, 18^. Actin filaments deliver mechanical support to the growth cone that enables force to generate movement. As such, protrusive activity (outgrowth) of an axon through a complex extracellular environment requires actin cytoskeletal remodeling. This highly regulated and complex process requires more than 100 accessory proteins to control the balance between actin filament elongation and branching. Rac GTPases act as master switches during actin cytoskeletal remodeling, acting upstream of both actin elongation drivers such as Ena/VASP and actin branching drivers such as the Arp2/3 complex^13^.

The Rac GTPase, Rac1, harbors several major functional domains that are highly conserved in metazoa: the guanine-binding domains, the membrane-targeting region, and Switch 1 and 2 regions. The guanine-binding domains bind guanosine diphosphate (GDP) or guanosine triphosphate (GTP), and the guanine-binding status determines whether Rac1 is inactive (GDP-bound) or active (GTP-bound). The membrane-targeting region of Rac1 is prenylated at specific amino acids to direct the protein to the plasma membrane, its site of action^19^. Finally, the Switch 1 and 2 regions are important for coordinating interactions that the Rac GTPase forms with regulatory and effector molecules. The Rac GTPase activity status can induce conformation changes in the Switch 1 and 2 regions that specify which regulatory or effector molecule it interacts with^20, 21^. Therefore, Rac GTPase activity modulates the output of this molecular switch. Two major types of upstream regulators control Rac GTPase activation status. Guanine nucleotide exchange factors (GEFs) exchange bound GDP for GTP, thereby activating Rac GTPases; and GTPase-activating proteins (GAPs) inactivate Rac GTPases by enhancing their intrinsic GTPase activity^22, 23^.

Here, we identified a genetic lesion, *ced-10(rp100)*, that causes axon guidance and outgrowth defects in the PVQ VNC interneurons in *C. elegans*. We demonstrate that *ced-10(rp100)* acts recessively and causes reduced GTP binding to CED-10 *in vitro*. The dramatic outgrowth defects observed in the *ced-10(rp100)* strain are caused by a single amino acid substitution (G30E) in the Switch 1 region of the CED-10 protein. In contrast, other viable mutations in CED-10, which have genetic lesions in the Switch 2 region (G60R) or the membrane-targeting region (V190G), exhibit PVQ axon guidance defects but do not show PVQ outgrowth defects. Further, CED-10(G30E) animals do not present canonical phenotypes caused by the G60R and V190G mutations - such as low brood size and the accumulation of apoptotic cell corpses. This posits that mutant CED-10(G30E) protein adversely affects specific downstream neuronal pathways while leaving non-neuronal functions intact. Our data also demonstrate that there are domain-specific CED-10 functions that can confer context-dependent roles during animal development.

In an unbiased genetic modifier screen, we identified that NAB-1 (Neurabin homolog) and SYD-1 (RhoGAP-like), a known NAB-1-interacting protein, inhibit PVQ outgrowth in animals with reduced CED-10 function. This suggests that NAB-1 and SYD-1 inhibit Rac GTPase activity to control axon outgrowth through GAP-dependent inhibition. Together, this study delineates regulatory functions of Rac GTPases and conceptualizes how different domains exhibit tissue-specific regulatory functions during development.

## RESULTS

### A Mutation in the CED-10/Rac1 Switch 1 Region Causes Defects in Axon Outgrowth and Guidance

During embryogenesis, the PVQL/R interneurons project axons from the tail into the ipsilateral fascicle of the ventral nerve cord (VNC) and terminate at the nerve ring in the head (Figure 1A). We identified a spontaneous mutation, called *rp100*, which causes over 80% penetrant defects in PVQ development (Figure 1A-B). Specifically, *rp100* causes the PVQ neurons to be inappropriately guided to the contralateral side of the ventral nerve cord and causes premature termination of the PVQ neurons (Figure 1A-D). Elevated temperature is known to place an added burden on axonal development, potentially due to accelerated development and/or perturbation of signaling pathways. Therefore, we examined how temperature effects PVQ development in *rp100* mutant animals at 20°C or 25°C. We found that the PVQ outgrowth defects of *rp100* mutant animals increased from ~35% to ~60% when raised at 25°C (Figure 1C), indicating that increased temperature enhances the detrimental effects of the *rp100* mutation on PVQ development. We wished to distinguish whether the PVQ axon outgrowth defects are an inherent deficiency in outgrowth or whether they are a secondary consequence of defective guidance. To examine this, we asked whether outgrowth defects are always accompanied with a guidance defect. We observed that ~20% of the outgrowth defects were not associated with a detectable guidance defect (Figure 1D), implying that the *rp100* mutation affects a fundamental process required for axon outgrowth. To determine if the PVQ axonal defects observed in *rp100* mutant animals are developmental or due to defective maintenance of neuronal architecture, we examined L1 larvae raised at 20°C and found that the total PVQ defects (88%) and PVQ outgrowth defects (37%) are comparable to those observed in L4 larvae (Figure 1B-C), indicating that *rp100* causes defects in axonal development.

**Figure 1.**
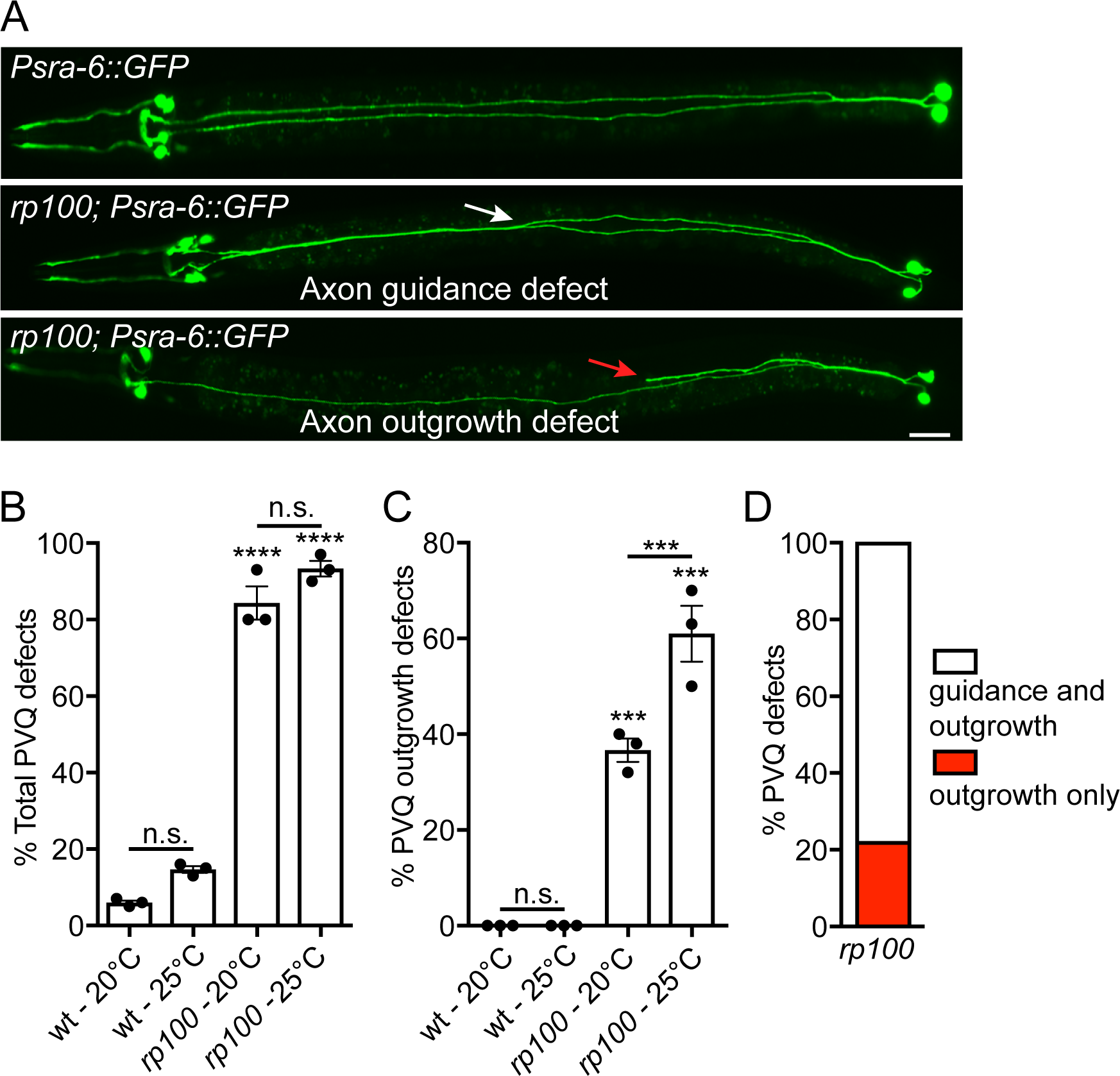
A spontaneous mutation causes severe defects in PVQ development. (A) Ventral views of animals expressing the transgene *Psra-6::gfp* to visualize the anatomy of the PVQ neurons (PVQL and PVQR). In wild type adult animals (upper image), the PVQ axons extend axons from the posterior into the ipsilateral side of the VNC. In *rp100* animals, PVQ axons inappropriately cross to the contralateral side of the VNC (middle image, white arrow) and/or exhibit premature termination of outgrowth (lower image, red arrow). Posterior to the right. Scale bar 20 μm. (B) The *rp100* genetic lesion causes highly penetrant defects in PVQ development. Wild type and *rp100* animals were incubated at 20°C or 25°C. Data are expressed as mean ±SD and statistical significance was assessed using t test. ^****^<0.0001, comparing wild type and *rp100* animals, n.s. not statistically significant. n=90 per strain, each dot represents independent scoring replicates. (C) The *rp100* genetic lesion causes highly penetrant, and temperature-sensitive, defects in PVQ axon outgrowth. Wild type and *rp100* animals were incubated at 20°C or 25°C. Data are expressed as mean ±SD and statistical significance was assessed using t test. ^***^<0.001, comparing wild type and *rp100* animals or *rp100* incubated at 20°C and 25°C, n.s. not statistically significant. n=90 per strain, each dot represents independent scoring replicates. (D) Phenotypic classification of PVQ outgrowth defects in *rp100* animals at 20°C. Approximately 80% of animals with outgrowth defects also have guidance defects (white bar) and 20% of outgrowth defects occur independently of a guidance defect (red bar). n=90, over three independent scorings.

To determine the molecular identity of the *rp100* genetic lesion we used a one-step whole-genome sequencing and SNP mapping method^24^. We found that *rp100* is a missense mutation, which results in an amino acid substitution from glycine (G) to glutamic acid (E) at position 30 in the Rac GTPase homolog CED-10 (Figures 2A and S1A). We confirmed that *ced-10(rp100)* causes PVQ outgrowth and guidance defects by rescuing both phenotypes through transgenic expression of a fosmid containing the *ced-10* locus (Figure 2B-C).

**Figure 2.**
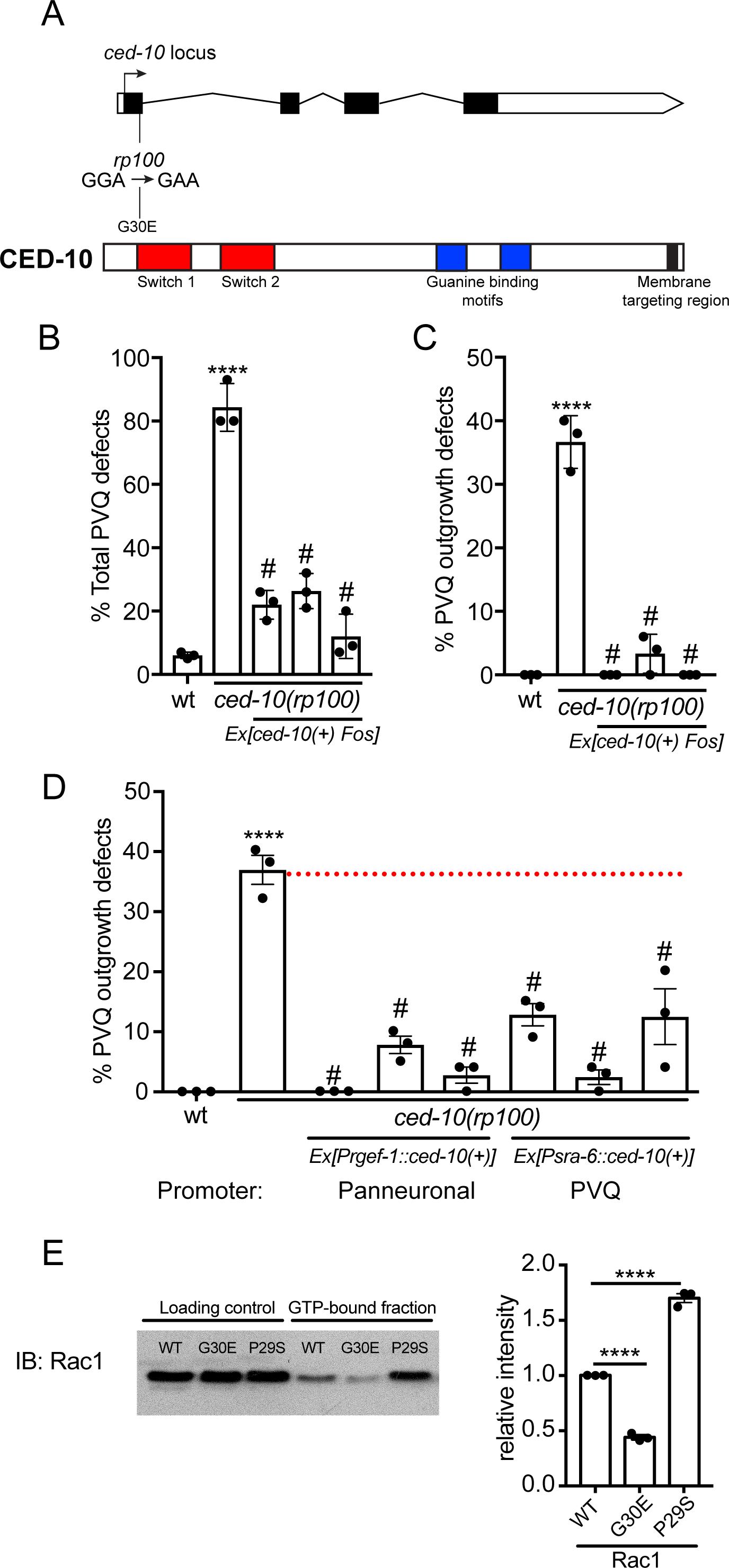
*rp100* is a mutation in the Switch 1 region of CED-10/Rac1. (A) Molecular identification of *rp100* as an allele of *ced-10*. *ced-10(rp100)* is a missense mutation in the first exon (GGA-GAA), causing a substitution of glycine at position 30 for a glutamic acid in the Switch 1 region of the Rac GTPase CED-10. The conserved domains within CED-10/Rac1 are marked as follows: Switch 1 and 2 regions (red boxes), guanine binding motifs (blue boxes) and membrane targeting region (black box). See Figure S1A for full sequence conservation between *C. elegans* CED-10 and human Rac1. (B-C) Transgenic expression of a fosmid containing the *ced-10* locus (WRM0639dH09) rescues both total PVQ defects (B) and outgrowth defects (C) of *ced-10(rp100)* animals. ^****^<0.0001, comparing wild type and *ced-10(rp100)* animals; #<0.0001 comparing the three transgenic rescue lines to *ced-10(rp100)*. n=90 per strain, each dot represents independent scoring replicates. (D) Transgenic expression of *ced-10* cDNA controlled by a pan-neuronal (*Prgef-1*) or PVQ-specific (*Psra-6*) promoter rescues PVQ outgrowth defects of *ced-10(rp100)* animals. Data are expressed as mean ±SD and statistical significance was assessed using one-way ANOVA, with Tukey’s multiple comparison test. ^****^<0.0001, comparing wild type and *ced-10(rp100)* animals; #<0.0001 comparing the three transgenic rescue lines to *ced-10(rp100)*. n=90 per strain, each dot represents independent scoring replicates. (E) *In vitro* pull-down of GTPγS-loaded recombinant Rac1 (wild type, G30E mutant or P29S mutant) with the GTPase-binding domain of PAK1. First three lanes = loading controls. Second three lanes = PAK1 binding assay to detect the GTP-bound fraction. Rac1(G30E) protein exhibits weaker PAK1 binding compared to wild type Rac1. The Rac1(P29S) gain-of-function mutant control shows excess PAK1 binding, as described previously^26^. Quantification of three independent experiments is shown in the graph. Data are expressed as mean ±SD and statistical significance was assessed using one-way ANOVA, with Tukey’s multiple comparison test. ^****^<0.0001, comparing each CED-10 mutant protein to wild type.

### CED-10 Acts Cell-Autonomously to Control PVQ Development

Rac GTPases are key regulators of cytoskeletal dynamics at the growth cone. As such, CED-10 is likely to act cell-autonomously to control PVQ axon outgrowth and guidance. To examine this, we expressed *ced-10* cDNA in the *ced-10(rp100)* strain using either the *rgef-1* promoter for pan-neuronal expression, or the *sra-6* promoter for PVQ-specific expression. We found that pan-neuronal and PVQ-specific *ced-10* expression robustly rescued PVQ outgrowth defects and partially rescued PVQ guidance defects (Figure 2D and Figure S1B). Incomplete rescue of PVQ guidance defects could be due to inappropriate timing or level of *ced-10* expression using these heterologous promoters. Nonetheless, the sub-optimal cell-autonomous rescue of PVQ defects prompted us to perform mosaic analysis to confirm our rescue data (Figure S1C). We transgenically expressed *ced-10* cDNA panneuronally and then identified rare mosaic animals that lost the extrachromosal rescuing array in the PVQ neurons. Here we found that *ced-10* expression in the PVQ neurons is required for PVQ axon outgrowth and guidance (Figure S1C). Therefore, we conclude that CED-10 acts cell autonomously to regulate PVQ axon outgrowth and guidance.

### *ced-10(rp100)* is a Hypomorphic Allele

The glycine affected by the G30E substitution is situated at the N-terminus of the Switch 1 region, a region fully conserved between CED-10 and Rac1 (Figures 2A and S1A). The Switch 1 region is known to coordinate interactions with upstream regulatory and downstream effector molecules^25^. Another mutation associated with the Switch 1 region, Rac1(P29S) causes gain-of-function effects to Rac GTPases (excessive GTP binding)^26^. Therefore, we used a combination of genetics and *in vitro* activity assays to ask whether 1) *ced-10(rp100)* acts as a gain- or loss-of-function mutation, 2) the G30E substitution affects the activity of the protein and 3) a known CED-10 gain-of-function mutation can cause PVQ axon outgrowth defects.

First, we examined whether *ced-10(rp100)* acts recessively by crossing a wild type chromosome into the homozygous mutant. We found that *ced-10(rp100)*/+ heterozygotes resemble wild type animals for both PVQ outgrowth and guidance, therefore the *rp100* mutation is recessive (Figure S2A-B). Next, we analysed PVQ outgrowth in the balanced *ced-10(tm597)* maternal-effect null allele. The majority of *ced-10(tm597)* animals derived from homozygous mothers die as embryos, however we identified L1 escapers, thereby enabling analysis of PVQ neuroanatomy (Figure S2C). We found that surviving *ced-10(tm597)* L1s exhibit ~60% outgrowth defects, indicating that defects in PVQ outgrowth is a *ced-10* loss-of-function phenotype (Figure S2C).

The activation status of Rac GTPases can affect their ability to regulate downstream signaling pathways during axon outgrowth^27^. We therefore tested whether the G30E mutation alters the ability of Rac1 to exchange GDP for GTP. To assay the ability of wild type and mutant Rac1 GTPases to perform nucleotide exchange, we *in vitro* loaded recombinant Rac1-GDP with a non-hydrolyzable GTP analog. Then, we used beads conjugated to p21-binding domain (PDB) of the Rac effector protein p21 activated kinase (PAK) to pull down the Rac1-GTP protein (Figure 2E). PAK-PDB binds with high specificity and affinity to Rac GTPases that are GTP-bound and not GDP-bound^28^. Our quantification of PAK-PDB-bound Rac1 revealed that, as previously reported, PAK-PDB pulled down a higher fraction of the Rac1(P29S) gainof-function mutant protein than wild type Rac1^26^ (Figure 2E). In contrast, a lower fraction of Rac1(G30E) was pulled down with PAK-PDB beads than wild-type Rac1, suggesting that less Rac1(G30E) protein is GTP-bound after *in vitro* loading compared to wild type Rac1 (Figure 2E).

Next, we used CRISPR Cas9 genome editing to introduce the fast cycling gain-of-function mutation (P29S) into the CED-10 protein and examined PVQ development^26, 29^. We found that animals expressing CED-10(P29S) exhibit 20% axon guidance defects and 0% axon outgrowth defects (n=90). Therefore, a known gain-of-function mutation does not have a strong deleterious effect on PVQ development. Together, these data support our genetic experiments showing that the *ced-10(rp100)* mutation causes a partial loss of CED-10 function.

### CED-10(G30E) Acts in Parallel with MIG-2 to Control PVQ Outgrowth

The GEF proteins UNC-73/Trio and TIAM-1 are known to positively regulate Rac GTPase activity in the *C. elegans* nervous system^27, 30^. We analysed PVQ development in *unc-73(e936)* and *tiam-1(ok772)* mutant animals and found that loss of *unc-73*, but not *tiam-1*, results in PVQ outgrowth defects (Table S1). Moreover, the *unc-73(e936); ced-10(rp100)* double mutant does not exhibit a synergistic effect on axon outgrowth (Table S1). These data indicate that UNC-73 is the major GEF for CED-10 during PVQ development. Modulating the affinity of GEFs for Rac proteins can fine-tune Rac activity. For example, the neuronal Navigator 1 protein NAV1 binds to Trio, which enhances the affinity of Trio for Rac1, a regulatory mechanism needed to control neurite outgrowth in mammals^31^. In agreement with this, we found that UNC-53, the *C. elegans* homolog of Navigator proteins, is also required for PVQ axon outgrowth (Table S1). Thus, it is likely that UNC-53 and UNC-73 act upstream of CED-10 to regulate axon outgrowth.

Losing the function of the Rac GTPase regulators UNC-73 and UNC-53 causes outgrowth defects of higher penetrance (55% and 88%, respectively) than observed for *ced-10(rp100)* (37%). This suggests that other Rac GTPases may be activated by UNC-73 to control PVQ outgrowth. Two possible candidates are RAC-2, which is nearly identical in sequence to CED-10, and MIG-2, which is functionally similar to mammalian RhoG^32^. We therefore examined PVQ development in *rac-2(ok326)* and *mig-2(mu28)* mutant animals. We found *rac-2(ok326)* animals are comparable to wild type. However, we observed extensive guidance defects (66%) and minimal outgrowth defects (4%) in the *mig-2(mu28)* null mutant (Table S1). To ask whether these Rac GTPases act in parallel to CED-10 to control PVQ outgrowth we performed double mutant analysis. We found that *ced-10(rp100); mig-2(mu28)* double mutant animals have highly penetrant defects in PVQ outgrowth (73%) and guidance (100%), whereas the *ced-10(rp100); rac-2(ok326)* is not significantly different from *ced-10(rp100)* (Table S1). In addition, *ced-10(rp100); mig-2(mu28)* mutant animals are severely uncoordinated, a phenotype not observed in either single mutant. Thus, our data indicate that, as reported previously, multiple neurodevelopmental decisions are redundantly regulated by these Rac GTPases^16, 30^.

A previous study showed that the SRGP-1 GTPase activating protein (GAP) negatively regulates CED-10 in the context of apoptotic cell corpse removal^33^. We asked whether SRGP-1 can also act as a GAP in the nervous system by examining PVQ development in the *srgp-1(gk3017); ced-10(rp100)* double mutant. We observed that loss of SRGP-1 function reduces the PVQ axon outgrowth defects observed in *ced-10(rp100)* animals (Table S1). Together, these data suggest that upstream activators of Rac GTPase activity, UNC-53, UNC-73 and SRGP-1, control PVQ axon outgrowth through parallel CED-10 and MIG-2 pathways.

### PVQ Outgrowth Defects are Specific to the CED-10(G30E) Mutation

Null mutations in *ced-10* cause embryonic lethality in *C. elegans^16^*. Therefore, hypomorphic alleles have traditionally been used to decipher biological functions for this Rac GTPase. *ced-10* alleles were originally isolated in genetic screens for mutants that are unable to execute apoptotic corpse engulfment^34, 35^. It was further shown that CED-10 acts in engulfing cells to coordinate cytoskeletal remodeling required for engulfment^36^. Two viable alleles isolated from these screens, *n3246* and *n1993*, affect the Switch 2 (G60R) and membrane targeting regions (V190G), respectively (Figure 3A). We asked whether these amino acid substitutions cause defects in CED-10 function that are important for PVQ development. We observed significant defects in PVQ axon guidance in the *n3246* (~80%) and *n1993* (~40%) *ced-10* alleles (Figure 3B). However, unlike the *rp100* allele, the *n3246* and *n1993* mutations do not cause defects in PVQ outgrowth (Figure 3C). We next performed extensive genetic analysis to delineate the impact of these *ced-10* hypomorphic mutations (*rp100*, *n3246* and *n1993*) on PVQ development (Figure S2A-B). First, we found that all three alleles are recessive, such that one wild type copy of *ced-10* is able to coordinate PVQ axon outgrowth and guidance (Figure S2A-B). We then generated transheterozygote combinations between the alleles. We noticed that all transheterozygotes exhibited 30-40% PVQ developmental defects, which is approximately half the penetrance observed in *rp100* or *n3246* homozygotes (Figure S2A). This indicates that none of the mutated CED-10 proteins are able to fully compensate for each other. In contrast, both the *n3246* and *n1993* alleles can fully compensate the PVQ outgrowth defects of *rp100* animals (Figure S2B). The *n1993/rp100* transheterozygote exhibits a residual ~10% penetrant defect in PVQ axon outgrowth. This phenotype is not significantly different from wild type, however, may suggest that a CED-10 protein with diminished membrane targeting cannot fully compensate for the defects caused by CED-10(G30E). Taken together, these data show that the G30E substitution in the *rp100* allele has a specific effect on PVQ axon outgrowth, not observed in other hypomorphic mutations in *ced-10*.

**Figure 3.**
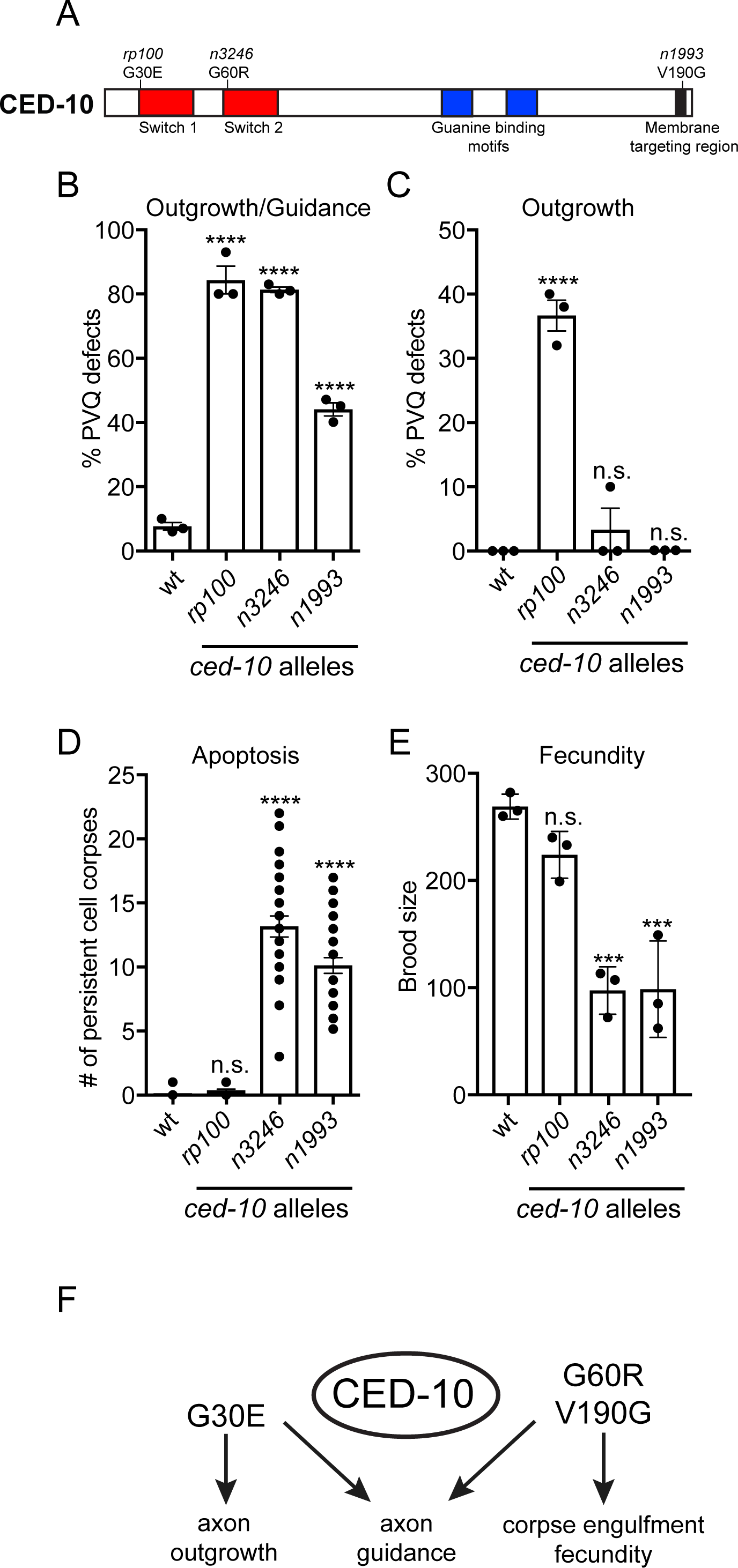
Context-specific effects of *ced-10* alleles. (A) Protein domain structure of CED-10 showing the hypomorphic alleles used in our analysis that affect specific domains. *rp100*(G30E): Switch 1 region; *n3246*(G60R): Switch 2 region and *n1993*(V190G): Membrane-targeting region. (B) All *ced-10* mutant alleles exhibit defects in PVQ development. Data are expressed as mean ±SD and statistical significance was assessed using one-way ANOVA, with Tukey’s multiple comparison test. ^****^<0.0001, comparing *ced-10* alleles to wild type animals. n=90 per strain, each dot represents independent scoring replicates. (C) Only *ced-10(rp100)*, and not the *n3246* and *n1993* alleles, exhibit PVQ outgrowth defects. Data are expressed as mean ±SD and statistical significance was assessed using one-way ANOVA, with Tukey’s multiple comparison test. ^****^<0.0001, n.s. not statistically significant comparing *ced-10* alleles to wild type animals. n=90 per strain, each dot represents independent scoring replicates. (D) Persistent apoptotic cell corpses are detected in the *n3246* and *n1993* alleles but not in *rp100* L1 animals. Data are expressed as mean ±SD and statistical significance was assessed using one-way ANOVA, with Tukey’s multiple comparison test. ^****^<0.0001, n.s. not statistically significant comparing wild type and *ced-10* alleles (n=30), over three independent scorings. (E) Brood size of *ced-10* hypomorphic alleles. *n3246* and *n1993* exhibit reduced brood size, whereas, *rp100* has a similar brood to wild type. Data are expressed as mean ±SD and statistical significance was assessed using one-way ANOVA, with Tukey’s multiple comparison test. ^***^<0.0001, n.s. not statistically significant comparing wild type and *ced-10* alleles n=30, each dot represents independent scoring replicates. (F) Summary of the effect of CED-10 hypomorphic mutations on PVQ axon outgrowth and guidance, apoptotic cell corpse engulfment and fecundity.

### CED-10(G30E) Does Not Disrupt Canonical CED-10 Functions

Thus far, we have shown that genetic lesions affecting specific domains, or residues, in the CED-10 protein can exhibit context-specific effects. We wanted to determine whether these differential phenotypic consequences are also observed for CED-10 function outside the nervous system. Therefore, we compared the effect of the *rp100*, *n3246* and *n1993* alleles in other well-characterized *ced-10* phenotypes - apoptotic corpse engulfment and fecundity (Figure 3D-E)^16, 36^. The number of persistent apoptotic cell corpses was counted in the heads of freshly hatched larval stage 1 (L1) animals harboring mutations in three *ced-10* alleles (Figure 3D). As previously reported, multiple persistent cell corpses were present in *n3246* and *n1993* mutant animals (Figure 3D)^36^. In stark contrast, we did not detect any persistent cell corpses in *rp100* mutant animals, as in wild type animals (Figure 3D). To ask whether *rp100* causes defects in apoptotic corpse engulfment at an earlier developmental stage, we performed a time-course analysis during embryogenesis (Figure S3). We found that there was a statistically significant increase in the number of persistent cell corpses at the 1.5-fold and 2-fold stages of embryogenesis between *ced-10(rp100)* and wild type embryos; however, these corpses were cleared by the time of hatching (Figure S3C). We next asked whether we could enhance this partial delay in corpse engulfment in the *ced-10(rp100)* mutant by removing genes that are important for the two convergent pathways that control apoptotic corpse engulfment (Figure S3D). Double mutants of *ced-10(rp100)* and the adaptor protein CED-6/GULP^37^, or the guanine nucleotide exchange complex subunit CED-12/ELMO^38^, had no significant effect on the number of persistent apoptotic corpses present at the L1 stage (Figure S3D). Taken together, the *ced-10(rp100)* mutation has minimal, if any, effect on apoptotic corpse engulfment, which defines it from the other known *ced-10* alleles.

We have shown that the molecular pathways dysregulated in *ced-10(rp100)* mutant animals do not adversely affect apoptotic corpse engulfment. In a reciprocal experiment, we asked whether signalling components through which CED-10 controls apoptotic cell recognition, engulfment and removal are required for PVQ axon outgrowth^39^. We examined the function of CED-1/LRP1/MEGF10, CED-2/CrkII and CED-5/Dock180, and found that these components of the apoptotic signaling pathway are dispensable for PVQ axon outgrowth (Figure S4A). We did find that *ced-5(tm1949)* mutant animals exhibit ~35% penetrant defects in PVQ guidance (Figure S4B), which suggests a function for CED-5 upstream of CED-10 in axonal development, as previously suggested^27^. These data show that the detrimental effect of the *ced-10(rp100)* mutation on PVQ axon outgrowth is not likely due to defective interpretations of signals from the core apoptotic corpse engulfment machinery.

CED-10 is also important for coordinating how cells change shape and move during embryogenesis^40, 41^. As a result, reduction of CED-10 function normally results in embryonic lethality and reduced brood size^42^. We therefore counted the broods of *rp100*, *n3246* and *n1993* mutant animals (Figure 3E). As previously reported, the *n3246* and *n1993* alleles cause a marked reduction in brood, whereas, the *rp100* allele generates a similar brood size to wild type animals (Figure 3E). Taken together, these data show that mutations in CED-10 have context-specific effects during *C. elegans* development, potentially due to differential requirements of regulatory and/or effector interactions with specific regions of CED-10 and/or CED-10 activity (Figure 3F).

### CED-10 Domain Mutations Have Neuron-Specific Effects During Development

Multiple decisions during neuronal development require CED-10 function^16, 35, 43^. To understand the broader impact of the *ced-10(rp100*) mutation on neurodevelopment compared to the *n3246* and *n1993* mutations, we crossed all three alleles into fluorescent reporter strains to permit visualization of neuronal development at single-neuron resolution (Table S2). We found that, in general, the membrane targeting-defective *n1993* allele had the weakest effect on neuronal development. However, the *n1993* allele did cause guidance defects in VNC neurons, left/right (L/R) choice of the VD motor neurons, and in the PLM and PVM neurons (Table S1). In contrast, the two Switch region alleles, *rp100* and *n3246*, had stronger and comparable effects on neuronal guidance, with a few notable exceptions (Table S1). In general, neurons that navigate the VNC are more strongly affected by *rp100* than *n3246*. For example, defects in PVP and HSN axon guidance, and AVG outgrowth exhibited higher penetrance in *rp100* than *n3246* (Table S1). In addition, the *rp100* allele exhibits higher penetrant defects in DD and VD motor neuron commissure guidance (Table S1). In contrast, PDE, AQR and mechanosensory neurons (ALM, AVM, PLM and PVM) development is more strongly affected by the *n3246* allele (Table S1). Taken together, these data uncover the domain-specific roles of CED-10 in *C. elegans* neuronal outgrowth and guidance.

### CED-10(G30E) Causes Defects in PVQ Outgrowth Through Specific Effector Molecules

Rac GTPases interact with multiple regulatory and effector molecules to control a vast array of biological processes. The effect of the CED-10(G30E) mutation on axon outgrowth could be due to a change in affinity with interacting molecules and/or altered activity or expression of its effector molecules. To determine the pathways through which defective CED-10(G30E) protein may cause PVQ outgrowth defects, we performed single and double mutant analysis with *ced-10(rp100)* and candidate genes that encode known Rac GTPase interactors or are known to regulate cytoskeletal remodelling at the growth cone (Table S1).

We first focused our analysis on known Rac GTPase interactors. RIN-1 is a VPS9 domain protein that interacts with GTP-bound CED-10 and controls neuronal guidance downstream of Slit-Robo signaling^44^. We found that RIN-1 is not required for PVQ development and that the penetrance of *ced-10(rp100)* PVQ defects is not affected by the *rin-1(gk431)* mutation (Table S1). Next, we examined the Lamellipodin homolog MIG-10, a major regulator of actin polymerisation to promote axon outgrowth^45, 46^. Surprisingly, *mig-10(ct41)* null mutant animals exhibit minimal PVQ axon outgrowth defects (2%), albeit with highly penetrant guidance defects (75%), and does not affect the PVQ outgrowth penetrance of *ced-10(rp100)* mutant animals (Table S1). This indicates that MIG-10/Lamellipodin is not a crucial regulator of PVQ axon outgrowth. The *C. elegans* abLIM homolog UNC-115 is an actin modulatory protein that functions to control growth cone filopodia formation^47^^-^^49^. We found that UNC-115 is also dispensable for PVQ axon outgrowth and *unc-115(ky275)* null mutant animals exhibit a low penetrant axon guidance defect (Table S1). The penetrance of PVQ axon guidance defects of *unc-115(ky275); ced-10(rp100)* animals is additive when compared to either single mutant suggesting that these factors regulate PVQ guidance in parallel (Table S1).

Next, we analysed the function of the p21-activated kinases (PAKs), which are known downstream Rac GTPase effectors that are important for controlling cytoskeletal dynamics. Dimerized PAKs interact with GTP-bound Rac proteins, which relieves PAK self-inhibition and permit kinase domain activation^50^. We analysed PVQ development using mutant alleles of the *pak-1*, *pak-2* and *max-2* genes (Table S1). We found that neither the *pak-2(ok332)* nor *max-2(ok1904)* mutants caused PVQ outgrowth defects. However, *max-2(ok1904)* animals do exhibit PVQ axon guidance defects, consistent with the finding that MAX-2 can regulate axon guidance independently of Rac proteins^51^. Furthermore, removing these PAK proteins had no effect on *ced-10(rp100)* PVQ axon outgrowth defects. In contrast, we found that two independent alleles of *pak-1* suppress the PVQ outgrowth defects of *ced-10(rp100)* mutant animals, without affecting the penetrance of PVQ guidance defects (Table S1). One interpretation of these data is that PAK-1 acts in a parallel pathway to CED-10 in the PVQ neurons and that loss of PAK-1 affects growth cone dynamics such that the *ced-10(rp100)*-induced defects are suppressed. Alternatively, our observations could mean that CED-10(G30E) protein alters PAK-1-regulated actin dynamics such that axon outgrowth is specifically disrupted.

### Neurabin Inhibits PVQ Outgrowth

To identify additional molecular components that facilitate the *ced-10(rp100)* PVQ axon outgrowth defects, we performed a forward genetic suppressor screen (Figure 4A). Approximately 60% of *ced-10(rp100)* animals exhibit PVQ outgrowth defects when cultivated at 25°C (Figure 1C). Therefore, we mutagenized the *ced-10(rp100); oyIs14* strain with ethyl methanesulfonate and screened for reduced PVQ outgrowth defects in the progeny of 1000 F2 animals cultivated at 25°C. Of the four independent alleles that significantly suppress *ced-10(rp100)* PVQ outgrowth defects, we focus on *rp117*, a recessive allele that reduces *ced-10(rp100)* PVQ outgrowth defects at 20°C from ~35% to ~10% (Figure 4). Using bulk segregant mapping^52^, we identified a premature ochre stop codon (TAT-TAA) in *nab-1*, which encodes the sole *C. elegans* ortholog of mammalian Neurabin (Figure 4B)^53, 54^. Using two, independent, *nab-1* deletion alleles, *gk164* and *ok943*, we confirmed that loss of *nab-1* supresses the *ced-10(rp100)* PVQ outgrowth defects (Figure 4B-C). Interestingly, loss of *nab-1* does not suppress PVQ axon guidance defects (Figure S5) suggesting a specific role for NAB-1 in outgrowth. In addition, we found *nab-1* is dispensable for correct PVQ development in a wild type background (Figure S5). To ask whether NAB-1 functions cell-autonomously to inhibit PVQ axon outgrowth in the *ced-10(rp100)* mutant background, we expressed *nab-1* cDNA using a PVQ specific promoter (Figure 4D). We found that PVQ-expressed *nab-1* cDNA reversed the suppression of PVQ axon outgrowth defects observed in the *ced-10(rp100)*; *nab-1(ok943)* double mutant (Figure 4D). This confirms that NAB-1 functions cell-autonomously in the PVQ neurons to control axon outgrowth in animals with reduced CED-10 activity.

**Figure 4.**
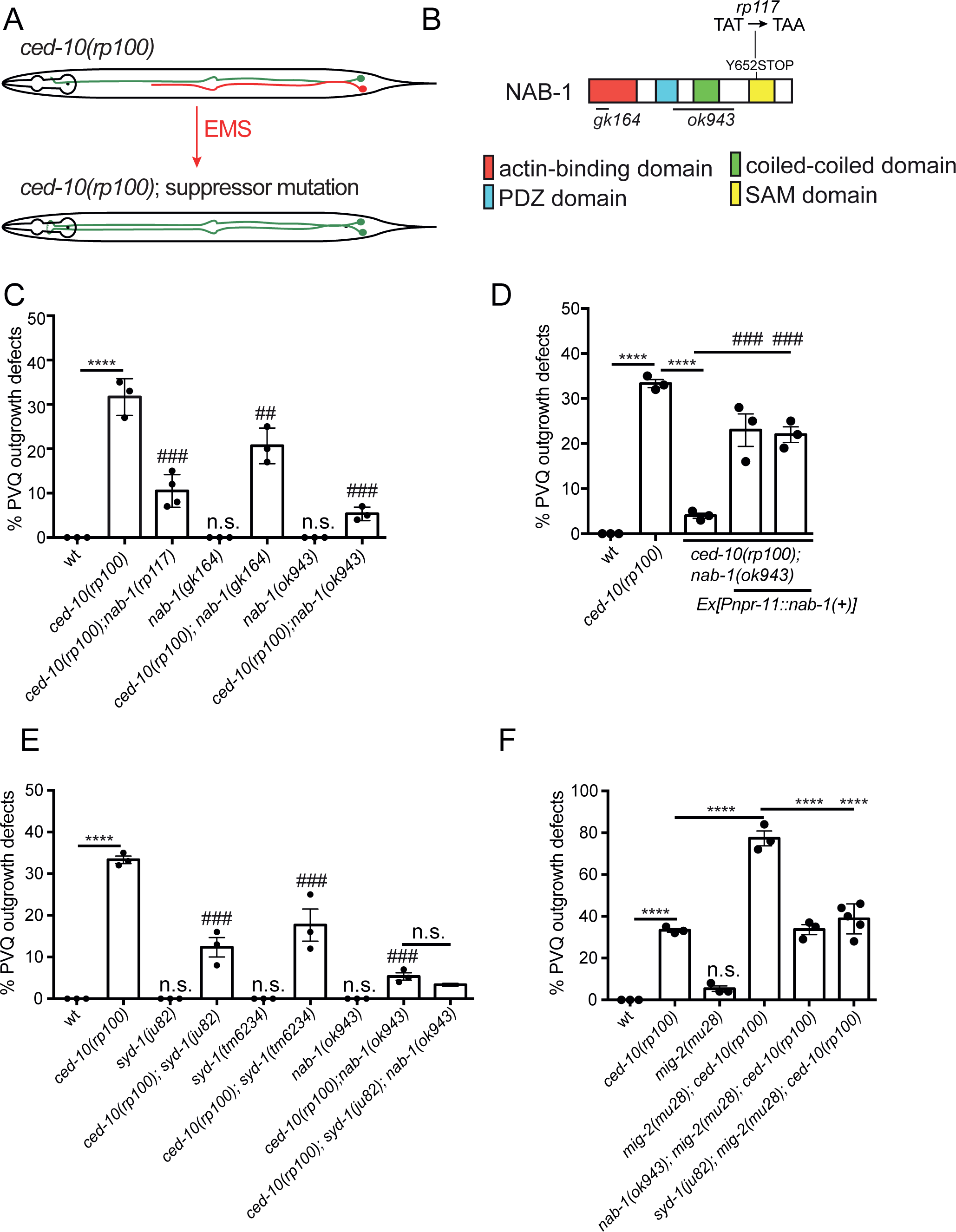
NAB-1 inhibits PVQ Axon Outgrowth. (A) Suppressor screen strategy (see main text). PVQL/R were marked by the *Psra-6::gfp* transgene. An outgrowth defect in the *ced-100(rp100)* strain is depicted by the red axon in the upper schematic. (B) Protein domain structure of *C. elegans* NAB-1. The allele obtained from the suppressor screen (*rp117*) and deletion strains (*gk164* and *ok943*) used in this study are indicated. (C) Loss-of-function alleles of *nab-1* (*rp117, gk164* and *ok943)* suppress the PVQ outgrowth defects of *ced-10(rp100)* animals. Data are expressed as mean ±SD and statistical significance was assessed using one-way ANOVA, with Tukey’s multiple comparison test. ^****^<0.0001, comparing wild type and *ced-10(rp100)* animals, ##=0.0006 and ###<0.0001, comparing compound mutants to *ced-10(rp100)*. n.s. not statistically significant when compared to wild type. n=90 per strain, each dot represents independent scoring replicates. (D) Transgenic expression of *nab-1* cDNA controlled by a PVQ-specific (*Pnpr-11*) promoter reverses the suppression of *ced-10(rp100)* PVQ outgrowth defects in *ced-10(rp100); nab-1(ok943)* animals. Data are expressed as mean ±SD and statistical significance was assessed using one-way ANOVA, with Tukey’s multiple comparison test. ^****^<0.0001, comparing wild type and *ced-10(rp100)* animals or *ced-10(rp100)* and *ced-10(rp100); nab-1(ok943)* animals; ###<0.0001 comparing the two transgenic rescue lines to *ced-10(rp100); nab-1(ok943)*. n=90 per strain, each dot represents independent scoring replicates. (E) Loss-of-function alleles of *syd-1* (*ju82* and *tm6234)* suppress the PVQ outgrowth defects of *ced-10(rp100)* animals. The *ced-10(rp100); syd-1(ju82); nab-1(ok943)* triple mutant has similar PVQ outgrowth defects as the *ced-10(rp100); nab-1(ok943)* double mutant. Data are expressed as mean ±SD and statistical significance was assessed using one-way ANOVA, with Tukey’s multiple comparison test. ^****^<0.0001, comparing wild type and *ced-10(rp100)* animals, ###<0.0001, comparing compound mutants to *ced-10(rp100)*. n.s. not statistically significant when compared to wild type or *ced-10(rp100); nab-1(ok943)* compared to *ced-10(rp100); syd-1(ju82); nab-1(ok943)*. n=90 per strain, each dot represents independent scoring replicates. (F) The highly penetrant PVQ outgrowth defects of *ced-10(rp100)*; *mig-2(mu28)* animals is suppressed by mutations in either *nab-1(ok943)* or *syd-1(ju82)*. Data are expressed as mean ±SD and statistical significance was assessed using one-way ANOVA, with Tukey’s multiple comparison test. ^****^<0.0001, n.s. not statistically significant when compared to wild type. n=90 per strain, each dot represents independent scoring replicates.

The functional domains of NAB-1 include an N-terminal actin-binding domain, a PDZ domain, a coiled-coil region and a C-terminal sterile alpha motif (SAM domain) with putative protein and RNA binding function^54^^-^^56^. Previous studies on NAB-1/Neurabin function have largely focussed on its role in synaptogenesis and synaptic function^53, 54, 57^. During *C. elegans* synaptogenesis, NAB-1 recruits the SYD-1 RhoGAP to filamentous actin^54^. SYD-1 itself has also been shown to interact with and repress MIG-2/RhoG to control HSN axon guidance^58^. We hypothesized that losing NAB-1, and potentially SYD-1 recruitment, would lead to Rac GTPase activation and subsequent suppression of the *ced-10(rp100)* axon outgrowth defects. To test this hypothesis, we generated compound mutants between *ced-10(rp100)* and two, independent, *syd-1* alleles (*ju82* and *tm6234*). We found that both *syd-1* alleles suppress *ced-10(rp100)* PVQ outgrowth defects (Figure 4E). Further, we found that removing both *nab-1* and *syd-1* did not further suppress *ced-10(rp100)* PVQ outgrowth defects, suggesting that these genes act in the same pathway to regulate neuronal outgrowth (Figure 4E).

Our data suggest that NAB-1 and SYD-1 are required to suppress Rac GTPase activity during PVQ axon outgrowth. We have shown that CED-10 and MIG-2 act redundantly in PVQ axon outgrowth (Table S1), and a previous study revealed that MIG-2 is negatively controlled by SYD-1^58^. Therefore, we hypothesized that loss of NAB-1 or SYD-1 activates MIG-2 and thereby promotes PVQ axon outgrowth. To test this hypothesis, we examined the effect of losing NAB-1 or SYD-1 in the *ced-10(rp100); mig-2(mu28)* double mutant, which exhibits highly penetrant defects in PVQ axon outgrowth (Figure 4F and Table S1). If NAB-1 and SYD-1 act through MIG-2, we would expect no suppression of PVQ axon outgrowth defects in *ced-10(rp100); mig-2(mu28)* animals. However, we found that PVQ axon outgrowth defects of *ced-10(rp100); mig-2(mu28)* animals are suppressed when *nab-1* or *syd-1* are absent (Figure 4F). As *mig-2(mu28)* is a null allele, these data suggest that NAB-1 and SYD-1 can regulate the activity of alternative GTPases, or may directly regulate CED-10 to control PVQ axon outgrowth.

## DISCUSSION

Axon outgrowth and guidance is coordinated by a plethora of extracellular signalling cues. Within the nascent axon, these cues are integrated through multiple, overlapping and non-overlapping, intracellular signaling cascades to ensure correct neuronal development. Rac GTPases are central to these process by acting as molecular switches to rapidly relay information through these signalling networks to enable appropriate axonal responses. In this study, we identified a single amino acid in the Switch 1 region of *C. elegans* CED-10/Rac1 that is crucial for regulating neuronal development. Focussing on the PVQ neurons, we found that a glycine to glutamic acid substitution at amino acid 30, CED-10(G30E), generates a CED-10 protein that causes severe axon outgrowth and guidance defects.

Our multiple lines of evidence show that the G30E substitution reduces CED-10 function. We have shown that *ced-10* null and *ced-10(rp100)* mutant animals exhibit similar PVQ axon outgrowth defects. In addition, PVQ axon outgrowth defects caused by *ced-10(rp100)* are fully recessive and are rescued by expressing wild type *ced-10* cDNA. We also demonstrate that animals harboring a known Rac1 hyperactive mutation (proline to serine substitution at position 29) in CED-10 did not exhibit PVQ axon outgrowth defects. Most importantly, using *in vitro* GTP activation assays we found that CED-10(G30E) is GTP activated ~50% less than wild type CED-10. The reduced affinity of CED-10(G30E) for GTP may result in diminished and/or atypical interactions between CED-10(G30E) and unknown effector molecule(s), thereby causing the PVQ axon outgrowth phenotype.

We found that the CED-10(G30E) mutant protein causes profound effects on PVQ outgrowth. However, unlike other CED-10 mutant proteins, CED-10(G30E) does not cause defects in apoptotic corpse engulfment or fecundity. This suggests that defects in cell behaviour caused by the CED-10(G30E) protein specifically affects nervous system development. It could also suggest that the activity of CED-10(G30E) is not sufficiently reduced to impinge upon apoptotic corpse engulfment or generation of progeny. Focusing our analysis on the nervous system, we found that the neuronal guidance defects caused by CED-10(G30E) are similar to another mutation affecting the Switch 2 region, CED-10(G60R). However, we only found PVQ axon outgrowth defects in CED-10(G30E) animals. To our knowledge, this is the first report where different regions of Rac GTPases have been shown to differentially regulate axon outgrowth and guidance. Our data therefore suggest that PVQ outgrowth is especially sensitive to CED-10(G30E)-defective signaling and is an ideal model to delineate previously unknown functions for the Switch 1 region of Rac GTPases in regulating cell behaviour. Further analysis of the actin cytoskeletal architecture in the PVQ neurons during embryogenesis - the time of PVQ outgrowth, is unfortunately not presently feasible, as the tools to observe PVQ-specific reporter proteins during embryogenesis are unavailable.

We performed a genetic suppressor screen to identify the regulators controlling CED-10-directed PVQ axon outgrowth. This approach revealed that loss of NAB-1, the *C. elegans* Neurabin homolog, suppresses *ced-10(rp100)* PVQ axon outgrowth defects. In *C. elegans*, NAB-1 regulates synaptogenesis and synaptic function by binding to F-actin and recruiting the synaptic active zone proteins SYD-1 (RhoGAP) and SYD-2 (liprin-α)*^53, 54^*. In mammals, Neurabin I was originally shown to promote neurite formation in primary rat hippocampal neurons^59^. Further, in a neuroblastoma cell line, Neurabin I was shown to directly interact with Rac3, an interaction that is required for Rac3 induction of neuritogenesis^60^. Neurabin II/Spinophilin also controls dendritic spine formation, with reduction of Neurabin II causing an increase in spine density and altered filopodia formation^61^. Subsequent work showed that both increases and decreases in Neurabin I levels affect neurite outgrowth, suggesting that changes in Neurabin expression can disrupt the balance in downstream signaling pathways^62^. Supporting this, Neurabin is known to interact with multiple Rho GTPase modulators including GEFs (Lfc and Kalirin) and a GAP (SYD-1)^54, 63, 64^. Therefore, as a scaffolding protein NAB-1 may either promote or inhibit neurite outgrowth depending on the whether it coordinates binding of a Rho GTPase activator (GEF) or inhibitor (GAP). Evidence for this was revealed in studies of rat cortical neurons, where overexpression of Neurabin I reduced Rac1 activation, whereas knockdown activated Rac1^62^. These data from mammalian systems support our findings that loss of NAB-1 promotes PVQ axon outgrowth in animals with diminished Rac GTPase activity. Further, our genetic data support previous biochemical studies where NAB-1 interacts with the Rho GAP SYD-1 to negatively regulate Rho GTPases^58^.

Taken together, our work reveals that a conserved amino acid (glycine 30) in the CED-10/Rac1 protein is important for the activation status of this Rac GTPase and for correct axon outgrowth of the PVQ neurons in *C. elegans*. Rac GTPase activation status is central for regulating downstream effector pathways, and we found that the scaffolding protein NAB-1 and RhoGAP SYD-1 likely enhance CED-10 activity and therefore modify CED-10 output. Intriguingly, we found that although reduced NAB-1 and SYD-1 function promoted axon outgrowth, it did not restore PVQ axon guidance. This supports the hypothesis that there are intracellular mechanisms in growth cones that are specifically regulated by effector molecule binding to Rac GTPases, which differentially promote/inhibit axon outgrowth and guidance. We propose that identifying the precise effector molecules and pathways controlled by CED-10 under different levels of activation will uncover novel insights into how correct axon outgrowth and guidance is achieved.

## METHODS

### Nematode Strains and Genetics

All *C. elegans* strains were maintained at 20°C on NGM plates seeded with *Escherichia coli* OP50 bacteria, unless otherwise stated^65^. Strains were generated using standard genetic procedures and are listed in Tables S3 and S4. All strains were backcrossed to N2 at least three times before scoring or generating compound mutants. Genotypes were confirmed using PCR genotyping or Sanger sequencing with primers listed in Table S5.

### Genetic mapping of *ced-10(rp100)*

We initially observed PVQ axon outgrowth and guidance defects in the following strain - *nDf67(mir-51 mir-53* deletion); *oyIs14^66^*. After ten backcrosses with N2 males the PVQ outgrowth defects disappeared. We re-examined the ninth backcrossed strain, performed a backcross and randomly selected 50 F2 progeny. We screened the F3 progeny and found 14 plates that exhibited the PVQ outgrowth phenotype. Two of those where heterozygote for the *nDf67* deletion, from which we singled animals and genotyped for loss of the *nDf67* deletion. These animals still exhibited the PVQ outgrowth phenotype. We named the mutant allele *rp100* and used the one-step whole-genome sequencing and SNP mapping strategy to map the genetic lesion^24^. Males of the Hawaiian strain CB4856 were crossed with *rp100; oyIs14* hermaphrodites. Ten F1 progeny carrying the *oyIs14* transgene were picked to individual plates and allowed to self-fertilize. F2 progeny carrying the *oyIs14* transgene were picked to 250 individual plates and allowed to self-fertilize. Approximately 20 F3 progeny from each of the 250 F2 plates were scored for PVQ axon outgrowth defects. 47 plates were homozygous for the phenotype causing mutation, based on the presence of PVQ outgrowth defects. Progeny of these animals were pooled and DNA was isolated. Pooled genomic DNA was sequenced using Ilumina sequencing and the resultant sequencing data was analyzed using the Galaxy platform. Our mapping identified a single lesion (GGA-GAA) at base-pair position 89 within exon 1 of the *ced-10* gene, which was independently confirmed by Sanger sequencing.

### Clonal EMS mutagenesis screen

Random mutations in *ced-10(rp100); oyIs14* animals were induced with ethyl methanesulfonate (EMS) following the modified version of a previously described protocol^65^. Since incubation at 25°C increases the penetrance of PVQ axon outgrowth defects caused by *ced-10(rp100)*, screening for suppressor mutations was conducted at 25°C. The germlines of *ced-10(rp100); oyls14* animals (L4 stage) were mutagenised using 50mM EMS (Sigma) in M9 buffer (22 mM KH_2_PO_4_, 42 mM Na_2_HPO_4_, 86 mM NaCl) for 4 hours (room temperature), after which they were transferred to standard OP50 NGM plates for 72 hours (20°C). F1s were allowed to self-fertilise on individual plates (20°C). Four F2s (L4 stage) were randomly picked from each F1 and separated into individual plates for incubation at 25°C. F3 populations with PVQ outgrowth defects lower than 20% (n=50) were selected and retested in subsequent generations for heritability of suppression.

### Identification of *nab-1* as a suppressor of *ced-10(rp100)*

Suppressor gene identification was carried out using whole genome sequencing and bulk-segregant mapping^52^. Candidate lines carrying suppressor mutations were backcrossed with *ced-10(rp100); oyls14* and 200 recombinant F2s from each cross were separated into individual plates for incubation at 25°C. Recombinant F2s homozygous for the suppressor mutation were identified by scoring for suppression of PVQ outgrowth defects of F3 offspring. 40 F2 plates exhibited the suppressor phenotype and worms were washed off with M9 buffer and pooled for DNA extraction with Gentra Puregene kit (Qiagen). Paired-end whole genome sequencing (WGS) with Illumina NextSeq500 was performed and the causative genetic lesion identified using Mutation Identification in Model Organism Genomes (MiMoDd) (version 0.1.7.3).

### *C. elegans* plasmid generation

Plasmid inserts were confirmed using Sanger sequencing prior to use.

#### RJP273 *sra-6^prom^::ced-10cDNAb*

*ced-10* cDNA isoform b was amplified from an oligodT amplified cDNA library using the oligos 5’-ttggctagcgtcgacggtacatgcaagcgatcaaatgtg-3’ and 5’-agatatcaataccatggtacttagagcaccgtacactt-3’ and ligated into a *sra-6^prom^* plasmid with a pPD49.26 backbone using *KpnI*.

#### RJP272 *rgef-1^prom^::ced-10cDNAb*

*ced-10* cDNAb from RJP273 was ligated downstream of a *rgef-1^prom^* in a pPD49.26 backbone using *Nhel-Spel.*

#### RJP296 *sra-6^prom^::GFP*

Sequence encoding GFP was ligated into a *sra-6^prom^* plasmid with a pPD49.26 backbone using *Nhel-Spel*.

#### RJP297 *sra-6^prom^::mCherry*

Sequence encoding mCherry protein was ligated into a *sra-6^prom^* plasmid with a pPD49.26 backbone using *Nhel-Spel*.

#### RJP370 *npr-11^prom^::nab-1*

*nab-1* cDNA was amplified from an oligodT amplified cDNA library using the oligos 5’- gctagcatgacaacggcttccgagc −3’ and 5’- ggtacctcacatgggaattgtgtgtgc-3’ and ligated into a *npr-11^prom^* plasmid with a pPD49.26 backbone using *Nhel-Kpnl*.

### Fluorescence microscopy

Animals were mounted on 5% agarose pads and immobilized using 50mM NaN_3_. Examination and imaging of neurons was performed using an automated fluorescence microscope (Zeiss, AXIO Imager M2) and ZEN software (version 3.1).

### Neuroanatomical observation and scoring

Neurons were scored, blinded to phenotype, in L4/young adult stages, unless otherwise stated. Scoring criteria for each neuron is detailed in Table S2. The following transgenes were used to enable visualization of specific neurons. PVQs: *oyIs14*, *hdIs26, rpEx1640*; Mechanosensory neurons: (ALMs, AVM, PVM and PLMs) *zdIs5;* HSNs: *rpEx6;* PDEs: *IqIs2*; PVPs: *hdIs26*; AVGs: *otIs182*; D-type motor neurons: *oxIs12*; AQR: *rpIs8*. All neuronal scoring was repeated in triplicates on independent days, n=75, except for *oxIs12* scoring, n=35.

### Microinjections and transgenic animals

Transgenic animals were generated as previously described^67^. Rescue plasmids were injected directly into *ced-10(rp100); oyIs14*. The WRM0639dH09 fosmid spanning the entire *ced-10* locus was injected at 1ng/μl. For pan-neuronal rescue, RJP272 (*rgef-1^prom^::ced-10cDNAb*) was injected at 10ng/μl. For cell-autonomous rescue, RJP273 (*sra-6^prom^::ced-10cDNAb*) was injected at 50ng/μl and RJP370 *npr-11^prom^::nab-1* at 10ng/μl. RJP370 (*npr-11^prom^::nab-1*) was injected *into nab-1(ok943); ced-10(rp100); oyIs14* animals at 20ng/μl. *myo-2^prom^::mCherry* was injected at 5ng/μl as a co-injection marker in all transgenic animals.

### Mosaic analysis

For mosaic analysis, transgenic animals were generated by injecting RJP272 (*rgef-1^prom^::ced-10cDNAb*) at 10ng/μl, RJP297 (*sra-6^prom^::mCherry*) at 10ng/μl and *myo-2^prom^::mCherry* at 5ng/μl into the *ced-10(rp100); oyIs14* strain. A transgenic line was selected that exhibited rescue of the PVQ axon outgrowth and guidance defects. Transgenic animals from this line were then scored for phenotypic rescue of the PVQ defects in the presence and absence of the rescuing extrachromosomal array in the PVQ neurons by detection of *mCherry* fluorescence.

### Brood size

For each replicate, 10 individual mid-L4 larvae were placed on separate plates. Every 24 hrs, each worm was moved to a new plate and the previous plate was left for 24 hrs prior to scoring to allow enough time for all the laid eggs to hatch. The experiment was conducted in 3 replicates on independent days.

### Scoring of apoptotic corpses

Mixed culture plates were washed thoroughly five times with M9 to remove all worms, leaving eggs behind. After 20 mins, freshly hatched L1 larvae were mounted on agarose and immediately scored by DIC optics for persistent cell corpses in the head (anterior to the posterior bulb of the pharynx).

### Bacterial expression plasmid generation

pGEX-4T-1-TEV-Rac1_1-177_ was obtained from Christina Lucato^68^. Point mutations in the Rac1 plasmids were generated using the Q5 site-directed mutagenesis kit (New England Biolabs) with the following oligonucleotides: pGEX-4T-1-TEV-Rac1(G30E): 5’-gcatttcctgaagaatatatccc-3’, 5’-attggttgtgtaactgatcag-3’and pGEX-4T-1-TEV-Rac1(P29S):5’-caatgcattttctggagaatatatc-3’,5’-gttgtgtaactgatcagtag-3’.

### Protein purification of wild type and mutant Rac1 (Residues 1-177)

pGEX-4T-1-TEV-Rac1 wild type and mutant plasmids were transformed into *Escherichia coli* BL21(DE3) CodonPlus cells and GST-Rac1 protein expression was induced by IPTG at 18°C overnight. Cells were pelleted and resuspended in lysis buffer (20 mM Tris, pH 8.0, 500 mM NaCl, 2 mM DTT, and 2 mM EDTA). Cells were lysed by sonication and cell debris was cleared by centrifugation. Soluble lysate was filtered at 0.8 μm. GST-Affinity purification was carried out with glutathione-Sepharose 4B resin (GE Healthcare) at 4°C for 90 min. The GST tag was cleaved with overnight incubation with TEV protease. Rac1 was eluted and further purified with size-exclusion chromatography on a HiLoad Superdex 75 16/60 column (GE Healthcare), equilibrated in SEC buffer (20 mM Tris, pH 8.0, 150 mM NaCl and 2 mM DTT).

### *in vitro* Rac1 activation assay

Purified wild type and mutant Rac1 proteins (500 μM) in SEC buffer was preloaded with 10 mM GDP and 5 mM EDTA, for 1 hour at room temperature. The reaction was terminated by adding 10 mM MgCl_2_ (final volume). EDTA and excess GDP is removed from GDP-loaded Rac1 with PD-10 desalting columns and equilibrated with 20 mM Tris, pH 8.0, 150 mM NaCl, 10 mM MgCl_2_, 2mM DTT. 30 μM Rac1-GDP was incubated with 500 μM GTPγS with or without 15 mM EDTA for 1 hour at room temperature and the reaction was terminated by adding 20 mM MgCl_2_ (final volume). 5 μg Rac1-nucleotide complexes were allowed to bind with 5 μg PAK-PDB beads (Cytoskeleton) in 500 μl of pulldown buffer (25mM Tris-HCl, pH 8.0, 40mM NaCl, 30mM MgCl_2_, 1 mM DTT, 1% (v/v) Igepal CA-630) for 1 hour at 4°C. Beads were washed three times with cold pulldown buffer. Active Rac1 bound to PAK beads were eluted in Lithium Dodecyl Sulfate loading buffer and subjected to SDS-PAGE. Eluted Rac1 protein was detected by western blotting and anti-Rac1 mouse monoclonal antibody (Cytoskeleton). Western quantification was carried out with ImageJ with intensities normalised to loading controls. Method adapted from^26^.

### Statistical analysis

Statistical analysis was carried out using Graphpad Prism 7 using one-way ANOVA with Tukey’s Multiple Comparison Test or t test, where applicable. Values are expressed as mean ±S.D. Values <0.05 were considered statistical significant.

## ACKNOWLEDGEMENTS

We thank members of Pocock laboratory and Brent Neumann for comments on the manuscript. Some strains used in this study were provided by the *Caenorhabditis* Genetics Center, which is funded by NIH Office of Research Infrastructure Programs (P40 OD010440), and by Shohei Mitani at the National Bioresource Project (Japan). We further thank Christina Lucato for help with production of recombinant proteins, Wolfgang Maier for expert advice on analysis of whole genome sequencing data and Cori Bargmann for the *npr-11* promoter. This work was supported by a grant from the European Research Council (ERC Starting Grant number 260807), Monash Biomedicine Discovery Fellowship, NHMRC Project Grant (GNT1105374), NHMRC Senior Research Fellowship (GNT1137645), and a Victorian Endowment for Science, Knowledge and Innovation Fellowship (VIF23) to R.P.

## AUTHOR CONTRIBUTIONS

S.N., S.D. and R.P. designed the research; S.N., S.D., W.C. and R.P. performed the research; S.N., S.D., W.C. and R.P. analysed data; and R.P. wrote the paper.

